# Framework for quality assessment of whole genome, cancer sequences

**DOI:** 10.1101/140921

**Authors:** Justin P. Whalley, Ivo Buchhalter, Esther Rheinbay, Keiran M. Raine, Kortine Kleinheinz, Miranda D. Stobbe, Johannes Werner, Sergi Beltran, Marta Gut, Daniel Huebschmann, Barbara Hutter, Dimitri Livitz, Marc Perry, Mara Rosenberg, Gordon Saksena, Jean-Rémi Trotta, Roland Eils, Jan Korbel, Daniela S. Gerhard, Peter Campbell, Gad Getz, Matthias Schlesner, Ivo G. Gut

## Abstract

Working with cancer whole genomes sequenced over a period of many years in different sequencing centres requires a validated framework to compare the quality of these sequences. The Pan-Cancer Analysis of Whole Genomes (PCAWG) of the International Cancer Genome Consortium (ICGC), a project a cohort of over 2800 donors provided us with the challenge of assessing the quality of the genome sequences. A non-redundant set of five quality control (QC) measurements were assembled and used to establish a star rating system. These QC measures reflect known differences in sequencing protocol and provide a guide to downstream analyses of these whole genome sequences. The resulting QC measures also allowed for exclusion samples of poor quality, providing researchers within PCAWG, and when the data is released for other researchers, a good idea of the sequencing quality. For a researcher wishing to apply the QC measures for their data we provide a Docker Container of the software used to calculate them. We believe that this is an effective framework of quality measures for whole genome, cancer sequences, which will be a useful addition to analytical pipelines, as it has to the PCAWG project.

## Introduction

Combining whole genome sequencing data from individual projects has many advantages: increased statistical power, the ability to extend hypotheses across several projects and the possibility of asking biological questions covering a wider range of phenomena. However when the genome sequencing data comes from different centres, was sequenced at different times and under different protocols, great care must be taken to ensure that the sequencing data is of comparable quality, to avoid drawing false conclusions. The Pan-Cancer Analysis of Whole Genomes (PCAWG) project provided us with a great opportunity to assemble, test and finalise which quality control measures are important for comparing the quality of whole genome, cancer sequences.

The PCAWG project assembled a cohort of 48 projects encompassed in the International Cancer Genome Consortium (ICGC)^1^ and The Cancer Genome Atlas (TCGA)^2^ of which we analysed 2959 cancer genomes (normal-tumour genome pairs) from 2830 donors. The size of the dataset and the diversity of the samples, representing many different cancers from varied populations, allow the exploration of many fundamental questions of cancer. There was inclusion criteria based on the sequencing platform (Illumina) and minimum sequencing depth. However there were 18 different sequencing centres involved and the sequencing was performed over a five-year time-span (2009-2014: a time period in which the sequencing methodology was evolving rapidly). To be able to perform analysis across the whole data set, it was necessary that the quality of the sequencing be carefully assessed.

There are advantages in a comprehensive set of quality measures. We will be able to exclude samples of low quality. This will save running downstream analyses, saving computational and the researchers’ time. Another advantage is for researchers in PCAWG studying driver mutations, we can provide a sanity check. If the driver mutation is only found in low quality samples, it may not be a good candidate, compared to if it is supported by high quality samples. As PCAWG will release the data for community to use, our quality measures will provide a guide to the quality of the whole genome sequences within. For researchers who wish to assess the quality of their whole genome cancer sequences, we have released our methods, in a Docker Container for easy implementation.

To develop a framework to determine the quality of samples, we use methods employed by the sequencing centres involved in PCAWG as well as results in the literature. TCGA marker papers (see references^3–5^ for examples from 2014-16) all include quality control (QC) measures such as depth of coverage, batch effects and contamination levels, calculated as part of the Firehose analysis infrastructure. Likewise a recent ICGC paper^6^ with samples sequenced from three different centres relied on similar QC measures computed by the Picard toolkit. Lu et al.^7^, carried out meta-analysis of exome data available from the TCGA for 12 cancer types which is similar, but not identical in scope, to the data set examined here. Their inclusion criteria were based on coverage depth and percentage of exome coverage for both the normal and tumour samples. Other cancer studies have also pointed to the importance of the percentage of the genome covered^8,9^ as well as error rates for each of the paired reads^10^ as QC measures.

Here we present the results of the work by sequencing centres and research groups involved in PCAWG to define important quality control measures, and how best to combine the results from these measures. Based on the PCAWG data we selected measures covering five important features to assess the quality of cancer genome sequences: mean coverage, evenness of coverage, somatic mutation calling coverage, paired reads mapping to different chromosomes and the ratio of difference in edits between paired reads, an edit being a base in the read which is different to the reference genome. These measurements we computed for both the normal and tumour samples. To summarise the five QC measures, we established a star rating system to cover the range of the highest quality cancer genomes, passing the thresholds set for each measurement, to those that had many sequencing quality issues.

## Result

All our analyses are based on the aligned sequences from the PCAWG core pipeline^11^. Within the aligned sequences we did not use duplicate reads, reads with a mapping quality of zero and ignored supplementary alignments (reads that map to more than one place in the genome). The first three quality control measures; mean coverage, evenness of coverage and somatic mutation calling coverage; are linked to different aspects of the coverage of the genomic sequence. The other two measures indicate discrepancies between the paired reads: mapping to different chromosomes and the ratio of edits between the paired reads compared to the reference genome. Finally we summarise these five measures into a star rating, for easy comparison of each of the sample pair’s quality.

**Mean Coverage** When deciding on what depth to sequence cancer genomes to, a trade off has to be made between the advantages of having a high coverage to the cost of sequencing. The higher the cancer genome is sequenced the greater the confidence in calling somatic events (see Alioto et al.^12^ for a comparison of somatic mutation calling at depths up to 300X). A precondition for the inclusion of a donor in the PCAWG study was the availability of a whole genome sequence of the normal and tumour with 25X coverage or greater. We found that a number of the projects submitting these genomes had calculated coverage differently. For standardization the mean number of reads covering each position in the genome was calculated, after low quality and duplicate reads were excluded so to not inflate the number of reads (see *Supplementary Methods* for exact methods used). As shown in *Supplementary Figure S1*, most commonly the normal samples were sequenced to around 30X, while there was a bimodal distribution for the tumour samples with maxima at 38X and 60X. To provide a meaningful guide to the quality of the genomes in PCAWG, we therefore set the thresholds for the mean coverage, after aligning, to 25X for normal samples and 30X for tumour samples. This resulted in 0.4% normal and 2.2% tumour samples not reaching these minimum criteria (*Supplementary Figure S1*).

**Evenness of Coverage** To confidently identify germline variants and somatic mutations, an even coverage across the target area^13^, in this case the entire genome, is ideal. For this QC measure we used two methods to test if the genome is evenly covered. One method is to calculate the ratio of the median coverage over the mean coverage (MoM). An evenly covered sequence should have a ratio of one, with the mean value the same as the median value, not skewed by very low or high coverage in certain regions. To decide within what range of values a sample should fall to be regarded as evenly covered, we used the whiskers of the boxplots in *Figure 1*, 1.5 × I.Q.R (interquartile range) of the data, which results in the range of 0.99 − 1.06 for a normal sample and the wider range of 0.92 − 1.09 for the tumour samples (*Supplementary Figure S2*).

**Figure 1:**
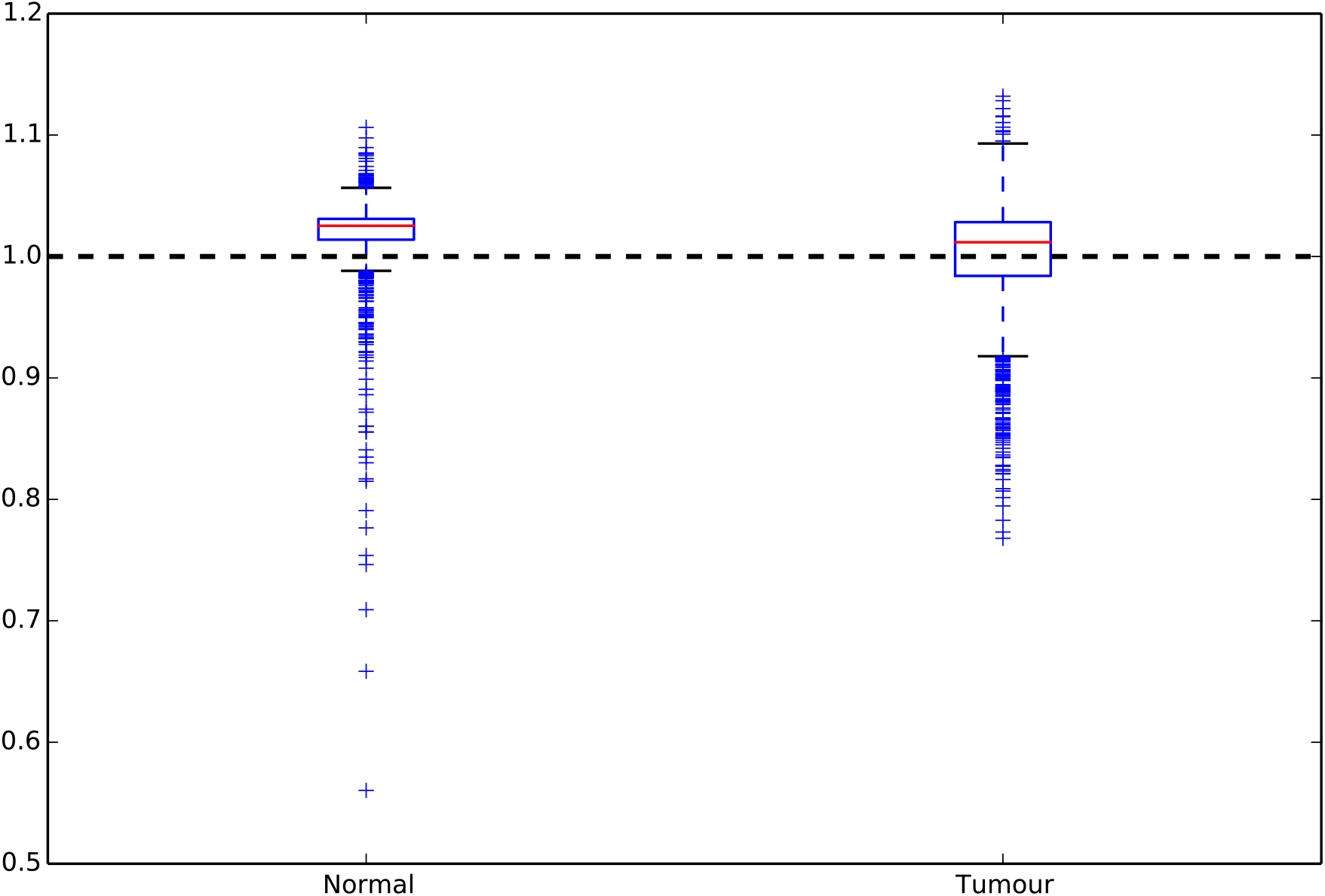
Distribution of the median coverage over mean coverage ratios for normal and tumour samples. The horizontal dashed bar at 1 represents the value of an evenly covered sample. As shown in the plot the tumour samples have a greater spread of values than the normal, we hypothesize this is to be expected as tumours are more likely to have deletions and structural rearrangements, which will lead to less evenly covered sequence. The whiskers on each of the boxplots (0.99−1.06 for the normal and 0.92−1.09 for the tumour) were taken as thresholds for this measure.

The second measure of evenness looks at the variation of the normalised coverage in ten kilobase genomic windows, after correction for GC-dependent coverage bias using the somatic CNV calling algorithm ACEseq^14^ (*Figure 2*). The main cloud, which corresponds to the main copy number state of the sample, is determined (as shown by the red dots in *Figure 2*). The remaining coverage variation is measured as full width at half maximum (FWHM) of the main cloud. This measure is insensitive to copy number aberrations and GC-dependent coverage bias. To determine the thresholds, 1000 WGS samples from different tumour types were used. We chose the thresholds based on clustering of these samples and subsequent visual inspection of the "best" samples that exceeded the threshold to see whether they are valid. Using these results the thresholds chosen are 0.205 for the normal and the more lenient 0.34 for the tumour, above which the sample would be regarded as having an uneven coverage (*Supplementary Figure S3*).

**Figure 2:**
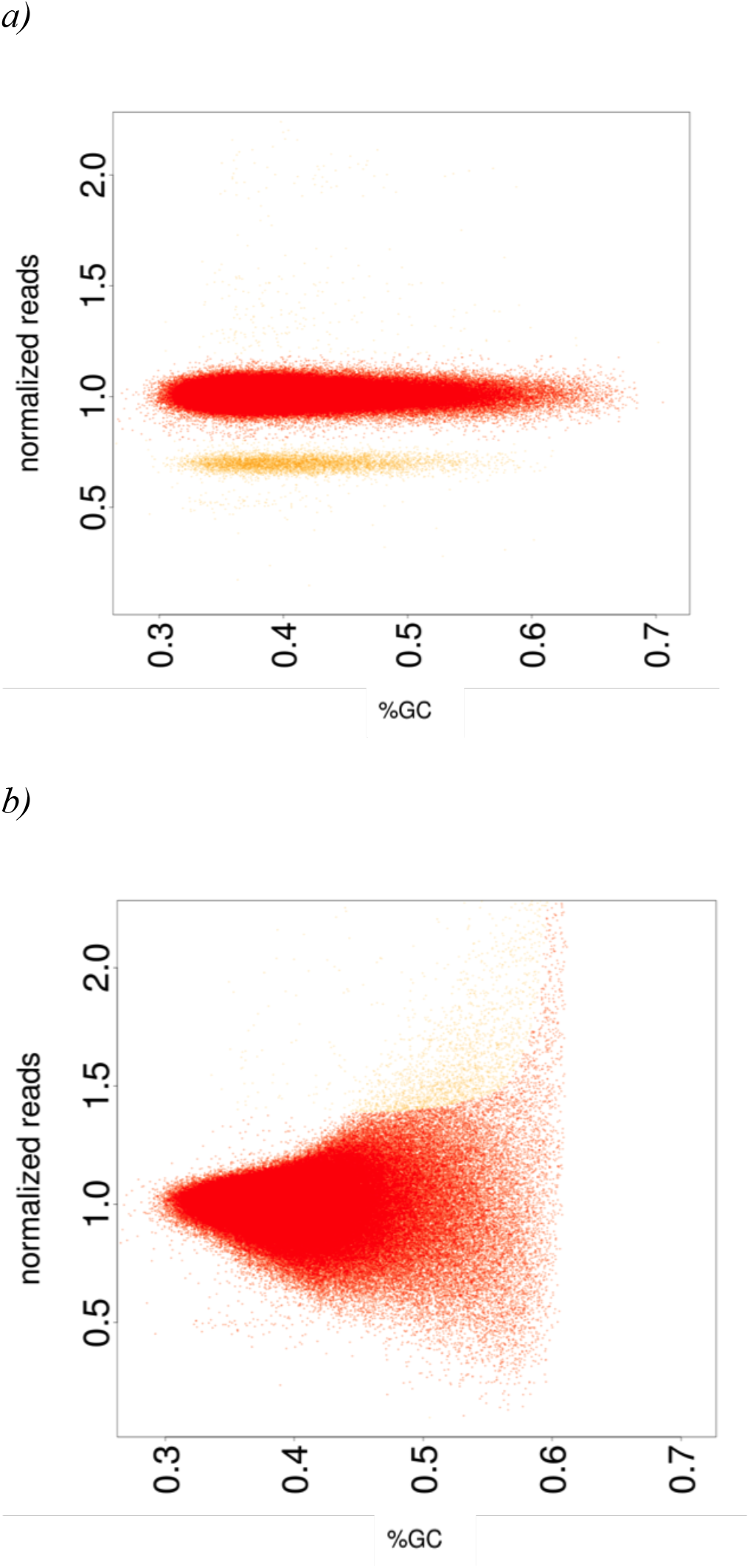
GC content versus the normalised coverage for evenly covered sample (a) and unevenly covered sample (b). The main cloud, corresponding to the main copy number state of the samples, is indicated in red. The yellow cloud represents a different copy number state of a copy number aberrant region. FWHM is calculated on the main copy number state.

For MoM coverage ratio and for FWHM, there is a greater range of values for the tumour samples than normal samples, potentially due to biologically reasons valid for tumours, for example large deletions could lead to a more unevenly covered sample. If the normal sample is unevenly covered, it is more likely due to a sequencing artefact. Hence, we are more stringent for the normal than the tumour samples.

The two evenness measures identify different samples as having uneven coverage (*Figure 3*). Spearman’s correlation coefficient for the two measures suggests that these measures are not correlated for the normal (ρ = 0.24) and tumour (ρ = −0.06) samples. FWHM is insensitive to GC bias, as the CNV caller corrects for this while MoM identifies other evenness outliers.

**Figure 3:**
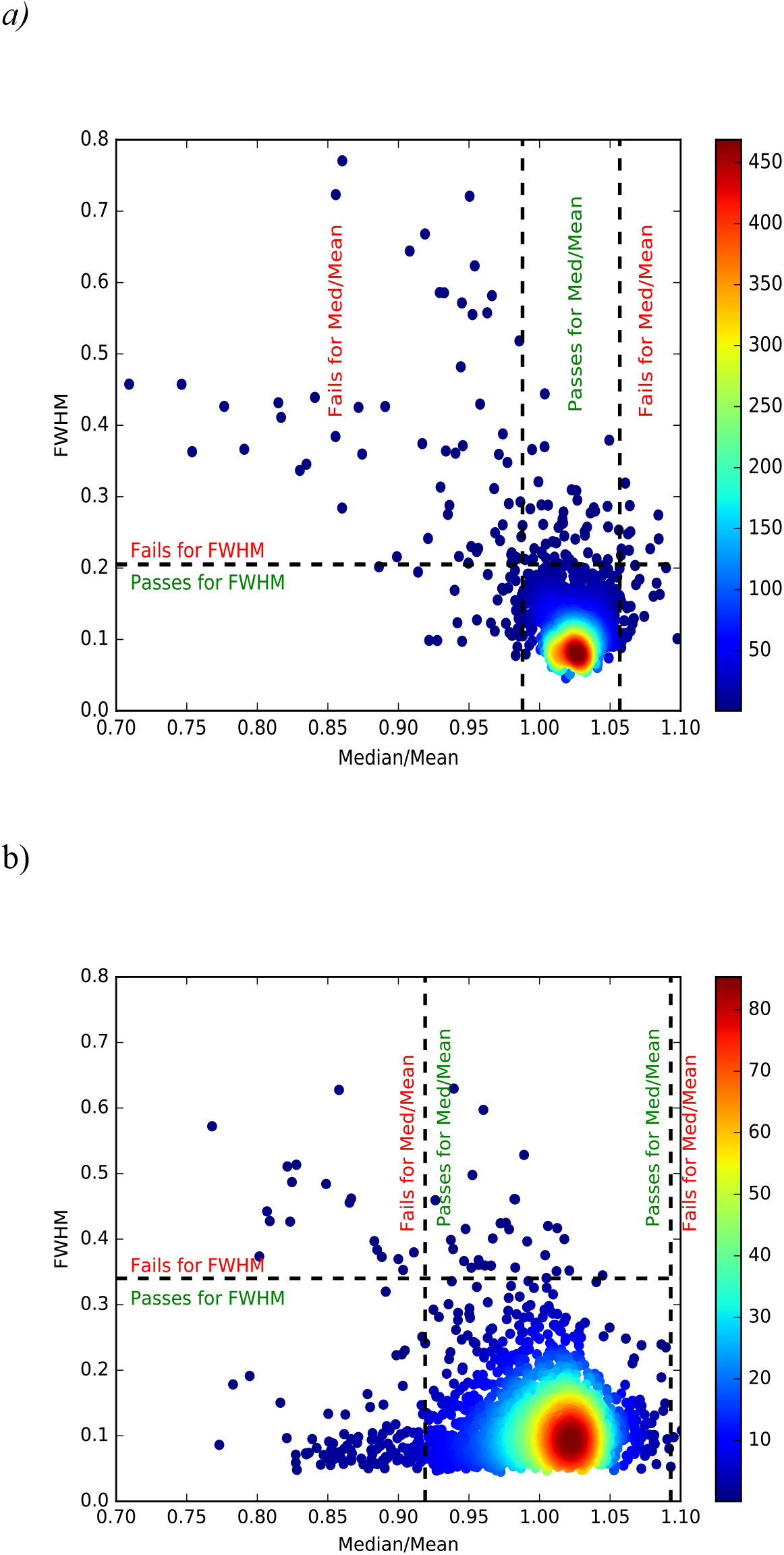
Density scatter plot comparing the two evenness of coverage measures for normal (a) and tumour (b). The number of samples overlapping is reflected by the colour at that point as shown by the legend. The dashed lines reflect the thresholds for the evenness measures. These graphs show that while there are certain samples both methods pick out as being unevenly covered, there are also samples picked out by one of the two.

The samples needs to be in the respective ranges of the MoM and below the thresholds for FWHM for the normal and the tumour to pass the evenness quality measure, of which 6.28% and 5.81% respectively of the samples were not.

**Somatic Mutation Calling Coverage** Having the depth of and evenness of coverage measured, our next QC measure looks at the effect of these at each base in the cancer genome (both the normal and the tumour sample). This measure gives a good summary of how much of the cancer genome is sufficiently covered to call a somatic mutation event. The somatic mutation caller MuTect^15^ calculates for each base in the genome, if it has sufficient coverage in both the normal and tumour sample (least fourteen reads are present in the tumour and eight reads in the matched normal sample). Based on those requirements, we had to establish the number of bases to consider the sample sufficiently covered. Ideally the threshold should be high enough to penalise the less well-sequenced samples, while not unduly penalising tumour samples that have had large deletions in the genome resulting in fewer bases to sequence. Taking into account the largest unambiguous mapping for a female donor (so not including the Y chromosome) would be 2,835,690,481 bases^16^, 2.6 gigabases would best suit these two needs. This results in 5.95% of normal-tumour pairs with fewer bases sufficiently covered, than this threshold (*Supplementary Figure S4*).

**Paired reads mapping to different chromosomes** The two reads from a read pair should represent the ends of a contiguous DNA sequence that depending on the insert size should be a given distance apart (for PCAWG between 200 and 1,000 bases). Paired reads mapping to different chromosomes can be due to a rearrangement. However an excess of reads mapping to different chromosomes points to a technical artefact. So deciding a threshold based on percentage of paired reads mapping to different chromosomes, we should not penalise sequences with biological causes of the paired reads mapping to different chromosomes (such as chromothripsis^17^, or more generally, interchromosomal rearrangements). We set the threshold to 3%, which even samples with confirmed high levels of rearrangements and chromothripsis do not exceed which in our experience, do not have more than 1% of paired reads mapping to different chromosomes. Of the normal sequences 14.5% exceed the threshold, as do 13.0% tumour sequences (*Supplementary Figure S5*). Interestingly there are more normal samples failing this measure, which cannot be explained by biological processes. A possible explanation may be that for lower quality samples in preparing libraries with PCR amplification causes an increase in two fragments of DNA from different parts of the genome being fused together, as has previously been noted^18^. Consequently, this translates to an increase in percentage of paired reads mapping to different chromosomes.

**Ratio of difference in edits between paired reads** Damage in sequencing runs has been linked to a global imbalance in edits (where the base in read is different compared to the reference) between read 1 and read 2 in paired end sequencing^19^. Therefore the ratio of the sum of edits between paired reads for a well-sequenced sample should be close to one. We adjudged samples with a two-fold ratio of edits between the paired reads, or greater, as having something gone wrong in the sequencing cycle resulting in lower data quality. Based on this threshold 4.66% and 4.49% normal and tumour samples failed respectively.

**Summary** The five quality measures were selected to provide minimal redundancy in flagging quality issues in normal/tumour paired genome sequences, which each measure reflects a facet of sequencing quality that other measures do not. *Figure 4* shows there is some overlap between certain measures, for example 75 sample pairs are penalised by both having a high percentage paired reads mapping to different chromosomes and uneven coverage. However a much higher number of samples penalised by one of these measures and not the other. Having defined these five, non-redundant QC measures our next step was to summarise them, to give an overall score for quality for the other researchers in PCAWG to use.

**Figure 4:**
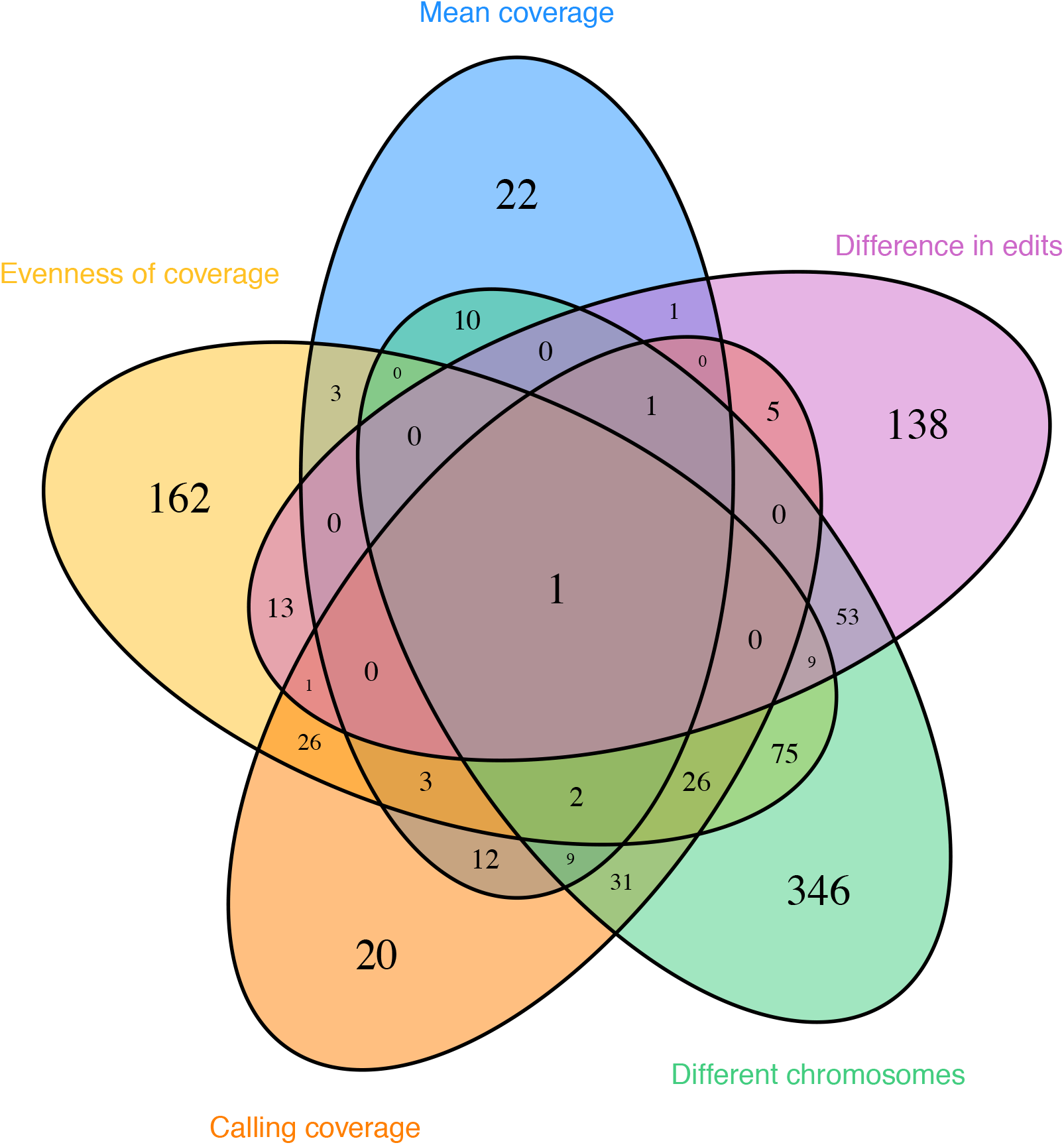
Venn diagram showing for which QC measure sample pairs were penalized for. The outside numbers show that each QC measures penalises a fair number of sample pairs uniquely. Looking at the overlaps between QC measures, while some measures are closer to each other than others, they all maintain a large degree of independence.

## Star rating system

We used the five quality measures to construct a star rating for each cancer genome (normal/tumour whole genome sequence). For each QC measure a star is awarded if both the normal and tumour sample pass the threshold. Half a star is awarded if only the normal passes the threshold for the respective QC measures. For somatic mutation calling coverage, a whole star is awarded for passing, none otherwise. The reasons for the extra weighting of the normal sample for the other four measures are that there is no biological reason for low quality in the normal sequence and a well-sequenced normal sample is important for calling somatic mutations.

Summing the stars earned for each of the five QC measure results in 66.4% of the normal/tumour sample pairs of the PCAWG being rated as 5 stars. Looking specifically at the different projects (*Figure 5*), a more nuanced picture is available. The quality does not seem to be biased by tissue type (*Supplementary Figure S7*) based on detailed molecular subtypes of the tumours in PCAWG^20^, the difference seems to be more at the project level. Unfortunately, there is only limited project metadata on when and which protocol was used to sequence the samples. Detailed metadata was available for 95 donors of the CLLE-ES project (concerning Chronic Lymphocytic Leukaemia), so it could be used as an example. Changes in protocol had an effect on the quality of the sequencing over the four years in which CLLE-ES samples were sequenced. For the CLLE-ES project, most notable was the change to a no PCR proband in 2012, which resulted in improvements to the measures of paired reads mapping to different chromosomes and evenness of coverage. This in turn resulted in a measurable change in somatic mutation calling coverage and improvement in star ratings (*Supplementary Figure S8*). We found similar results for a subset of 348 samples sequenced at the Broad Institute (*Supplementary Figure S9*), which had metadata recorded in CGHub^21^ about the time and instruments used to sequence. We hypothesise that this will be true for other projects as well.

**Figure 5:**
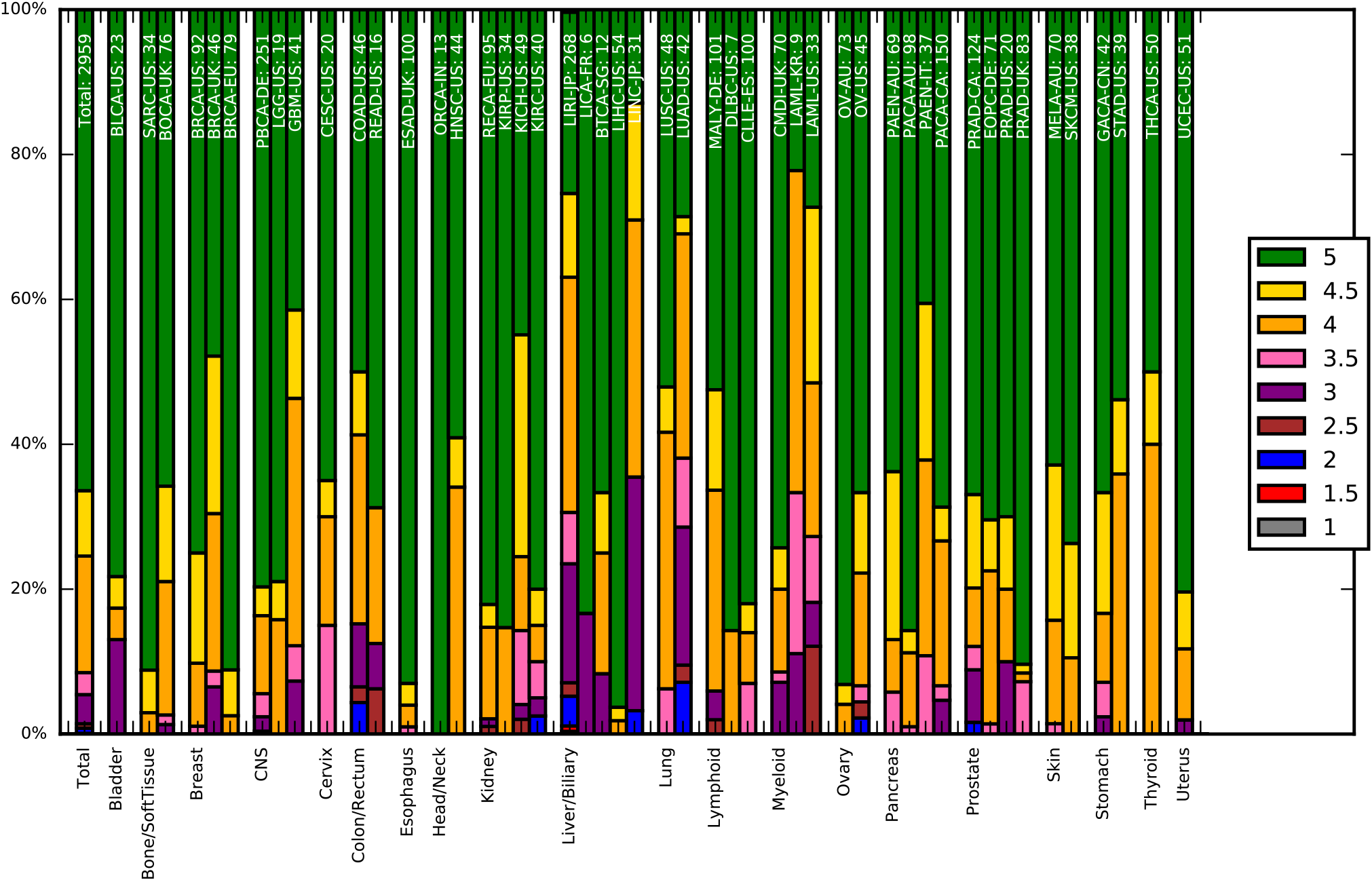
Distribution of the star ratings for the PCAWG genomes, grouped by tissue type (as labelled along the x-axis), and then project. The project name and number of samples in the project are labelled at the top of the bar. The colour of the bar reflects what percentage of samples in the project have that star rating (corresponding to the legend). The bar on the far left shows the results for all samples. The plot demonstrates the varying quality of different projects - differences we believe come from when the genome was sequenced and the sequencing protocol used.

Having calculated the star rating for the sequences, it was interesting to see how our QC measures relate to the calling of somatic single nucleotide variants (SNVs)^11^, somatic insertion and deletions (indels)^11^ and somatic structural variants (SVs)^22^ in PCAWG. An advantage of using these PCAWG datasets is that four callers were used for each. Looking at the proportion of calls, which all four callers supported, gives us a good idea how the quality of sequencing influences the identification of unambiguous somatic mutations. While the proportion of calls supporting the four callers varies greatly by sample, we find that the samples with four stars or more tended to have higher proportions than samples with less than four stars for SNVs, indels and SVs (with p-values of ~10^−5^, ~10^−5^, ~10^−18^ respectively, using the Mann-Whitney-U test, also see *Figure 6*).

**Figure 6a:**
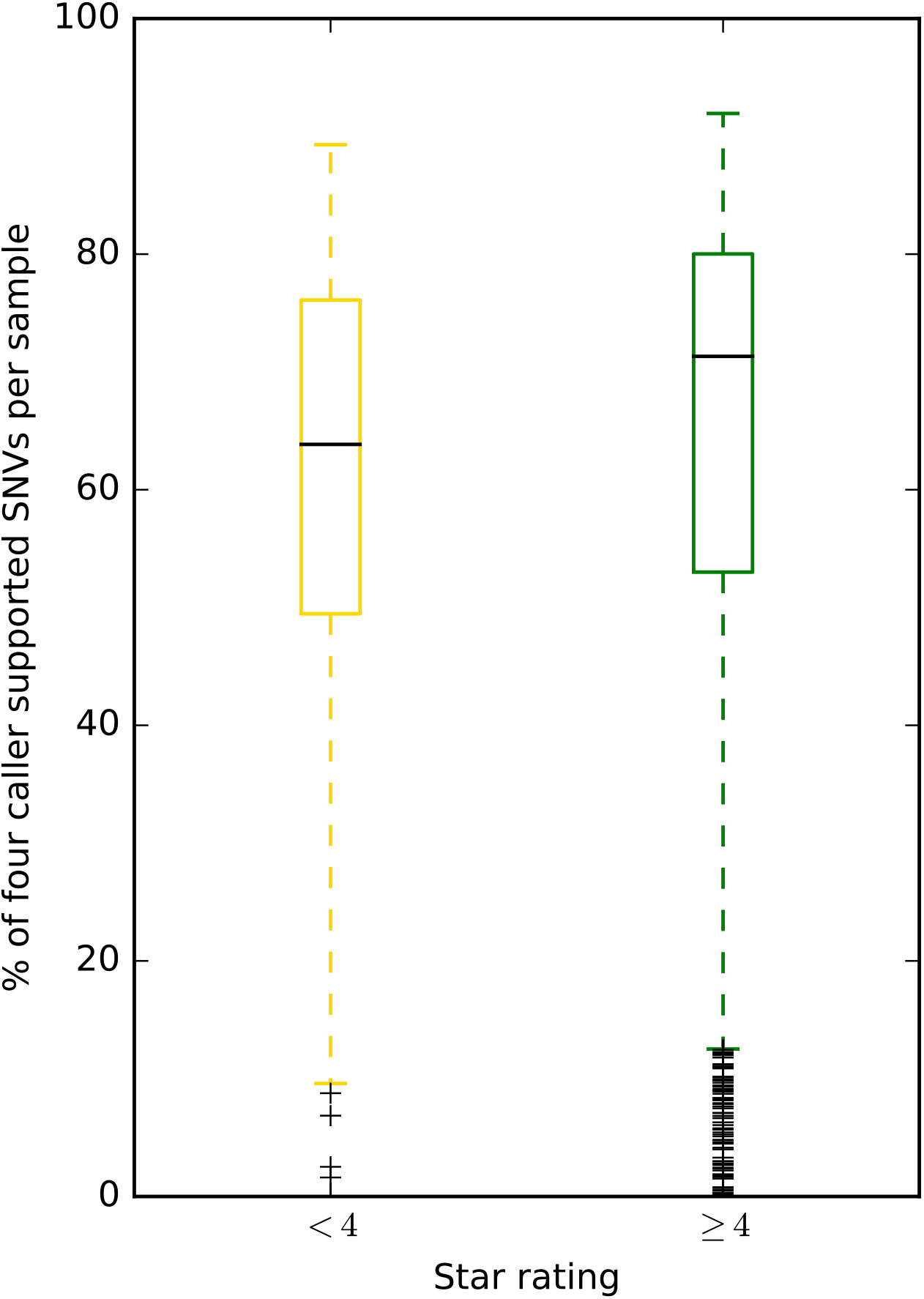
Samples with four stars or greater tend to have a higher the proportion of somatic single nucleotide variants (SNV) calls supported by four callers than samples with fewer than four stars. This is significant using the Mann-Whitney U test, with p-value ~ 10^−5^.

**Figure 6b:**
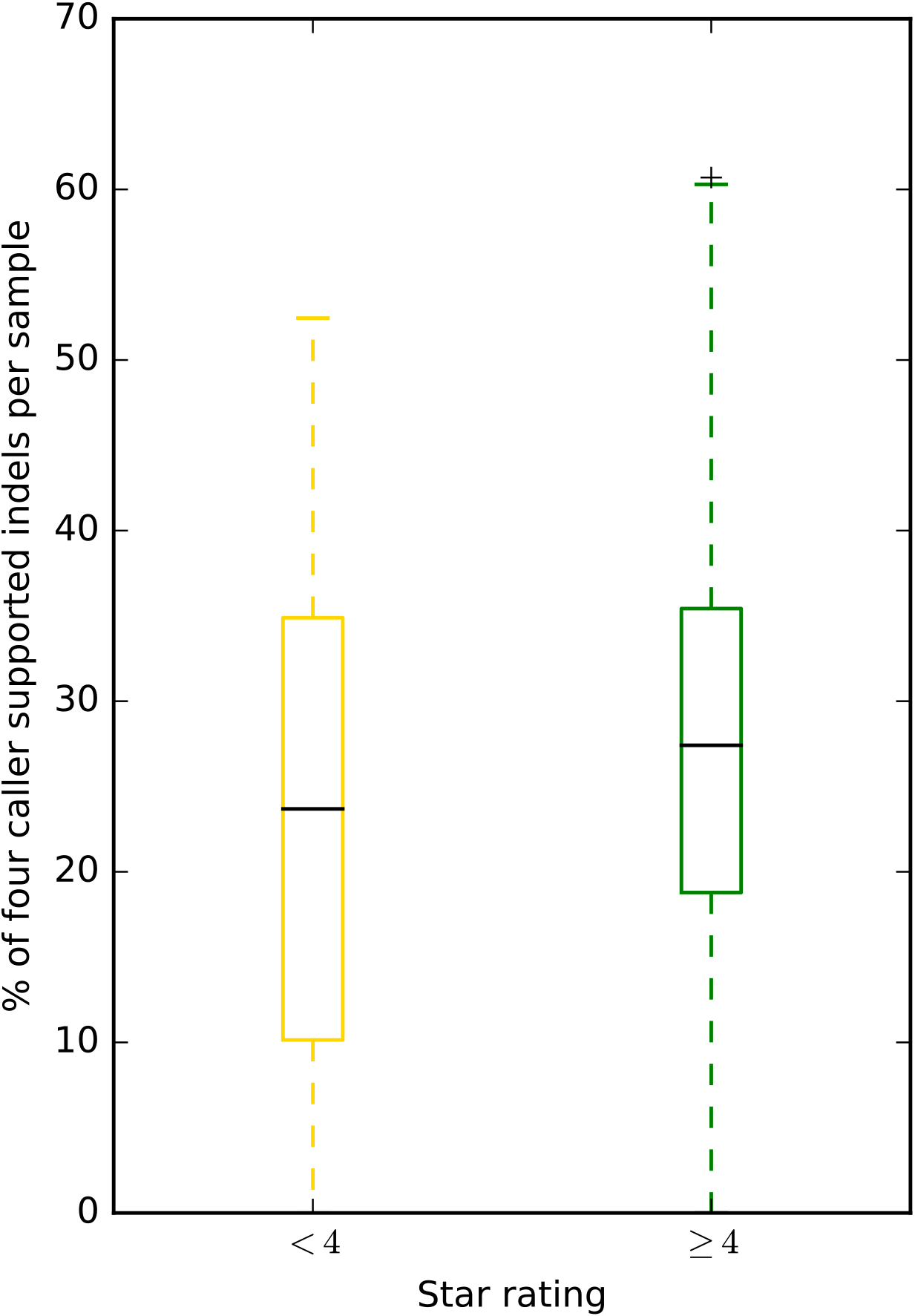
Samples with four stars or greater tend to have a higher the proportion of somatic insertion and deletion (indel) calls supported by four callers than samples with fewer than four stars. This is significant using the Mann-Whitney U test, with p-value ~ 10^−5^.

**Figure 6c:**
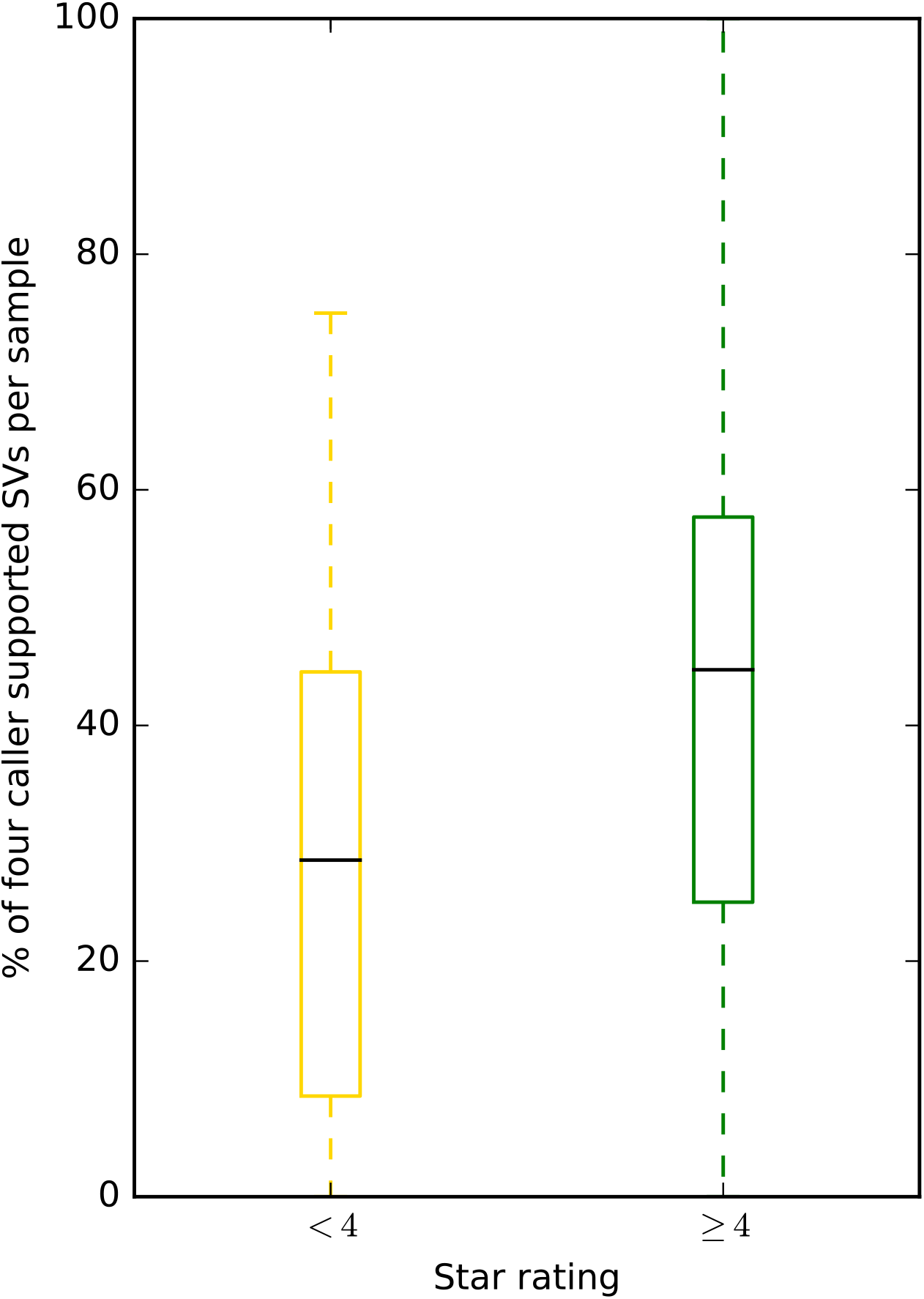
Samples with four stars or greater tend to have a higher the proportion of somatic structural variant (SV) calls supported by four callers than samples with fewer than four stars. This is significant using the Mann-Whitney U test, with p-value ~ 10^−8^.

Taking this analysis further we used linear regression models to further analyse the relation between the proportion of calls supported by four callers and the QC measures (see *Supplementary Tables S1-S3*). The results show that, an increasing percentage of paired reads mapping to different chromosomes in tumour samples, has a negative effect on the proportion of calls supported by four callers for SNVs, indels and SVs. For SNVs an increasing mean coverage in tumours has a significant positive effect on the proportion of calls supported by four callers. While for indels there is a significant negative effect on the proportion of calls supported by four callers by increasing unevenness (as measured by FWHM) in tumours. As in indels, the unevenness effect is also true in SVs as well as significant negative effects by increasing percentage of paired reads mapping to different chromosomes in normal samples and ratio of difference in edits between paired reads in tumour samples.

The results from this analysis suggest quality of sequencing, measured by our star rating, does have a measurable effect on the downstream analyses. As our QC measures reflect different aspects of sequencing quality, they also have varying levels of importance in using these sequences in the calling of SNVs, indels and SVs.

## Discussion

The established star rating system allows grading the normal and tumour sample sequences by quality in absence of information on how sequencing was carried out, what protocols were used and what problems may have occurred during the sequencing process. The system is not designed to be all encompassing, instead using a small amount of computational resources and time (compared to the actual aligning of the sequences), we get a good snapshot of the quality of the normal-tumour sample pair sequences on which to call somatic mutations. Likewise having graded the cancer genomes with our five-star system, we do not intend researchers to necessarily exclude the lower ranked cancer genomes, just to be wary of any conclusions based solely on the lower scoring genomes.

With our star rating system, we sent several samples in PCAWG to the exclusion list due to their poor performance in one of the QC measures. Due to the timing, this did not prevent the downstream analyses being performed. Though anecdotally it would have saved 55 days computational runtime for our one star sample. For all samples that remained, the QC star rating was embedded in the header of the variant call format files for use of the researchers within PCAWG, and when the data is released, to all researchers.

For those projects in PCAWG, which we had metadata, we found that sequencing quality has definitely improved over the time period 2009-2014 in which the samples sequenced. Our results for the CLLE-ES project suggest that in part a protocol change to PCR-free methods improved sequencing, as in line with best practices from a recent benchmarking exercise^12^.

Another advantage of our quality control is the link to the downstream analyses. In aggregate, the higher the quality of the sequences, had a higher proportion of the somatic SNVs, indels, SVs identified, by all the callers for each type of somatic mutation. These results suggest overall that higher quality sequence will identify the true positive somatic mutations with higher probability. Our data would suggest that when pre-amplification of DNA is needed for WGS, for example DNA isolated from formalin fixed, paraffin embedded tissue, the star rating system will be helpful when the variants and mutations are interpreted.

We believe that our method can be adapted for similar projects that look to use whole genome sequences from a variety of sources. The thresholds we used based on our experience and applied to this dataset of 2959 cancer genomes can also be used as guide to quality of sequences. It is worth noting that they represent a trade-off of being severe enough to penalise poor quality while not discriminating against samples with valid biological causes. We also would recommend using our methods to ascertain the quality before downstream analyses by other groups. To enable others to use our approach, there is a Docker Container, which can be accessed at https://github.com/eilslabs/PanCanQC. We provide a framework for quality assessment, which opens the door to do large-scale meta-analysis in a more robust framework.

## Acknowledgements

The authors would like to thank Jennifer Jennings and her colleagues at the Ontario Institute for Cancer Research (OICR) for their help in the administration of this working group.

JPW, MDS, SB, MG, JT and IGG are supported by the Ministerio de Economía, Industria y Competitividad and European Regional Development Fund (MINECO/FEDER BIO2015-71792-P), the Instituto de Salud Carlos III (ISCIII) and the Generalitat de Catalunya. In addition we have received funding from ELIXIR-EXCELERATE (EC H2020 #676559) and RD-Connect (EC FP7/2007-2013 #305444).

The work done by IB, KK, JW, DH, BH, RE and MS was supported by the BMBF-funded Heidelberg Center for Human Bioinformatics (HD-HuB) within the German Network for Bioinformatics Infrastructure (de.NBI) (#031A537A, #031A537C) and the BMBF-funded German ICGC-projects (ICGC-PedBrain: 109252 (German Cancer Aid), 01KU1201A,B; ICGC-MMML: 01KU1002B and ICGC-DE-MINING: 01KU1505E).

ER, DL, MR, GS and GG would like to acknowledge G.G. MGH startup package and Broad funds.

KMR and PC are members of the Cancer Genome Project supported by a Wellcome Trust grant (098051).

## Author contributions

JPW, IB, ER, KMR, KK, MDS and JW wrote the manuscript, helped develop and apply the methods and analysed the results.

SB, MG, DH, BH, DL, MP, MR, GS and JT contributed to the development of the methods.

RE, JK, DSG, PC, GG, MS and IGG provided project supervision; through feedback and the reviewing of the work done, as well as editing of the manuscript.

IB and JW constructed the Docker Container with code contributions from KK and KMR.

PCAWG-Tech and PCAWG-Network provided the data, metadata and the framework for this research.

